# Electric field based dosing for TMS

**DOI:** 10.1101/2023.07.31.551253

**Authors:** Ole Numssen, Philipp Kuhnke, Konstantin Weise, Gesa Hartwigsen

## Abstract

**Abstract:** Transcranial magnetic stimulation (TMS) is an invaluable non-invasive brain stimulation (NIBS) technique to modulate cortical activity and behavior, but high within- and between-participant variability limits its efficacy and reliability. Here, we explore the potential of electric field (e-field) based TMS dosing to reduce its variability and discuss current challenges as well as future pathways. In contrast to previous dosing approaches, e-field dosing better matches the stimulation strength across cortical areas, both within and across individuals. Challenges include methodological uncertainties of the e-field simulation, target definitions, and comparability of different stimulation thresholds across cortical areas and NIBS protocols. Despite these challenges, e-field dosing promises to substantially improve NIBS applications in neuroscientific research and personalized medicine.

**Outstanding Questions Box:** Outstanding Questions

- Does the cortical threshold for effective stimulation differ between primary regions and higher-level association areas? How large is the impact of cytoarchitectonic differences between regions on a stimulation threshold?
- Do cortical stimulation thresholds differ across individuals? Are thresholds stable within an individual across the lifespan? What are the physiological factors influencing these thresholds?
- Can a cortical stimulation threshold measured with single-pulse TMS be transferred to repetitive TMS protocols for the study of cognition?
- How does the cortical stimulation threshold interact with the current brain state?

**Graphical abstract:** 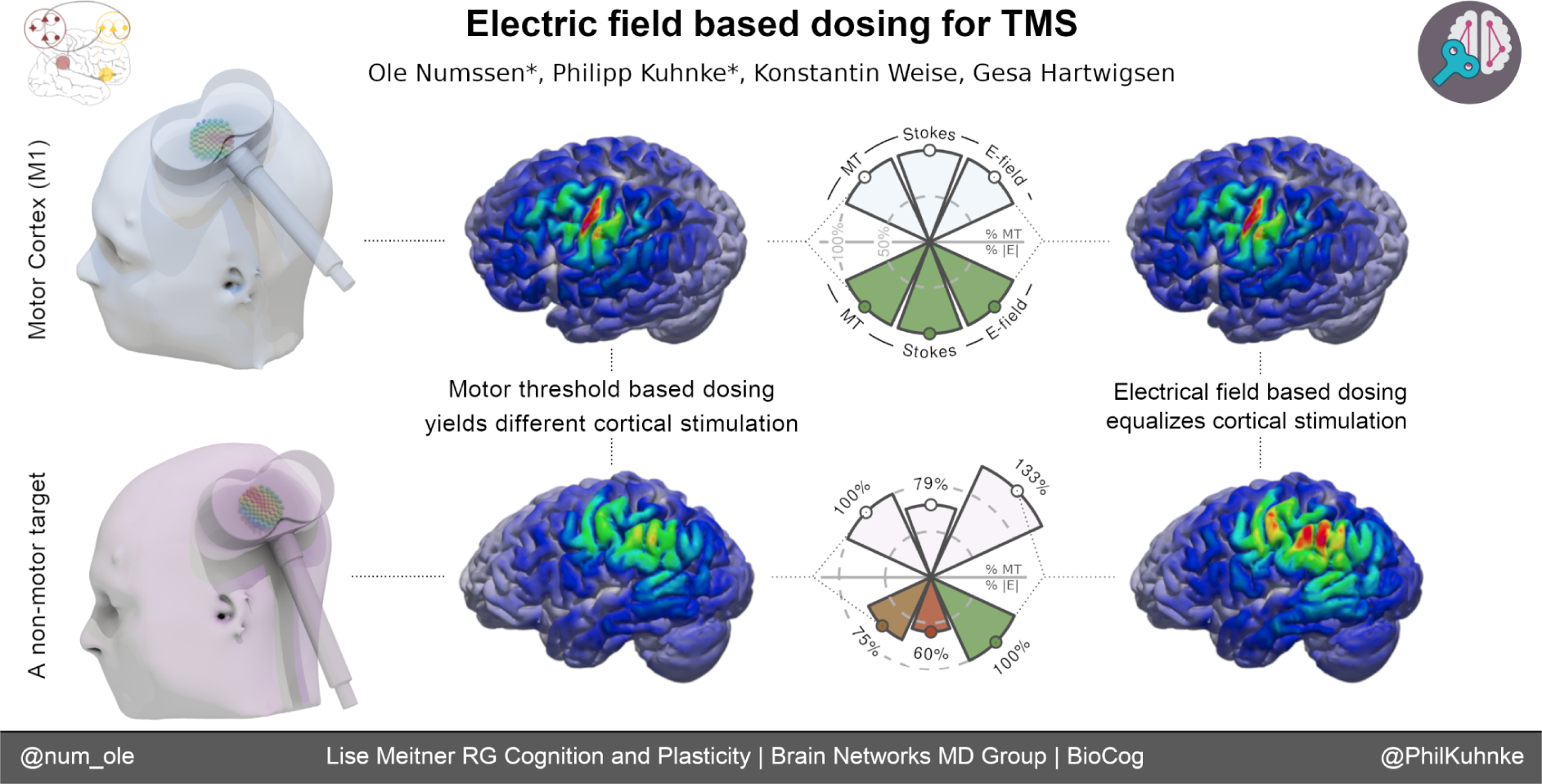

---

Non-invasive brain stimulation (NIBS) techniques, such as transcranial magnetic stimulation (TMS; Pascual-Leone et al., 2000), have emerged as invaluable tools for modulating brain activity in both healthy individuals (Walsh and Cowey, 2000) and psychiatric patients (Burt et al., 2002). Notably, TMS has received FDA approval as a therapeutic intervention for several disorders (e.g. O’Reardon et al., 2010). However, high variability of the stimulation effects within and across individuals limits strong conclusions about structure-function relationships (Numssen, van der Burght, et al., 2023; Caulfield et al., 2022). Likewise, the effect sizes in TMS studies are often small (Beynel et al., 2019). Consequently, there is an ongoing debate about the validity and reliability of different TMS protocols in research and treatment settings (e.g. Hartwigsen et al., 2015; Sandrini et al., 2011). Recent studies emphasize that differences in the individual responsiveness to NIBS strongly affect the outcomes and thus explain large parts of the observed variance (e.g., Hamada et al., 2013). One key factor determining NIBS effects, both for primary sensory-motor regions (Sasaki et al., 2018) and higher association areas (Lee et al., 2021), is the stimulation strength. Throughout this Perspective, we utilize the term *dosage* to refer to the strength of the TMS-induced cortical stimulation, targeted towards researchers and clinicians applying conventional TMS with available hard- and software. A broader view of potential influence factors is provided in the *Remaining challenges of e-field-based dosing* section. We also refer the reader to Peterchev et al. (2012) for an in-depth discussion of dosing parameters apart from the local stimulation strength. Currently, the gold standard for dosing across the brain is based on the individual motor cortex excitability and quantified via the motor threshold (MT) (Turi et al., 2021).

Here, we identify shortcomings of the MT-based dosing approach across various stimulation targets and suggest an alternative strategy. To this end, we present experimental data directly comparing different dosing approaches in the same set of individuals. Critically, standard MT-based dosing strategies fail to consider the actual level of stimulation of the cortical target due to their focus on the *stimulator* intensity (e.g., 50% maximum stimulator output; MSO). In contrast, we highlight the potential of TMS dosing based on the *cortical stimulation* itself, quantified via the induced electric field (Caulfield et al., 2021a; Weise & Numssen et al., 2022b; Dannhauer et al., 2022; Kuhnke & Numssen et al., 2023). Throughout this Perspective, *stimulator intensity* refers to the intensity that is set at the stimulator device (e.g., 50% MSO). In contrast, *stimulation strength* refers to the cortical stimulation exposure, quantified by the e-field strength (e.g., 100 V/m). It is essential to note that the TMS-induced e-field shows limited focality, rendering the exclusive stimulation of a single cortical location, without any off-target stimulation, impossible (Kuhnke et al., 2020). In our prior discussions (Numssen & van der Burght et al., 2023), we explored this limitation of TMS and other NIBS approaches and presented strategies to address these constraints. Here, we shift our focus to the singular cortical target—a spatially delimited region within the cortical gray matter. This specific region is the intended focal point for effective stimulation, guided by the presumed importance of this cortical structure in relation to a distinct functional domain. Consequently, we introduce the term “TMS-induced effect” to broadly characterize a cortical modulation that results in measurable effects on a designated outcome variable (see Hartwigsen & Silvanto (2022) for an in-depth discussion). This term encapsulates the observable impact stemming from the targeted cortical region, reflecting our focus on a localized, functionally relevant dosing approach.

Recent methodological advances have enabled the computational simulation of TMS-induced e-fields (e.g. SimNIBS, Puonti et al., 2020, Thielscher et al., 2015; ROAST, Huang et al., 2019), yielding the foundation for a biophysiologically informed dosing strategy. Calibrating the cortical stimulation exposure to the individual participant and brain region removes a critical variance source of TMS studies: the intra- and inter-individual variability in cortical stimulation exposure due to anatomical differences (Kuhnke & Numssen et al., 2023; Caulfield et al., 2021b). This Perspective utilizes single- and group-level analyses of cortical stimulation from TMS to identify shortcomings of dosing approaches that do not take the cortical stimulation into account. We reason that our e-field based dosing approach may also inform other transcranial electric and magnetic NIBS approaches that rely on induced electric fields (Peterchev et al., 2012), such as transcranial direct/alternating current stimulation (Jiang et al., 2020; Kasten et al., 2019; Laakso et al., 2019; Wischnewski et al., 2021), or temporal interference stimulation (Esmaeilpour et al., 2021). Although stimulation principles differ (e.g. supra-vs. sub-threshold stimulation), it was shown that differences in individual morphology affect the cortical stimulation exposure to a similar or even higher degree (Weise et al., 2022a).

While e-field based dosing represents an important step towards more reliable and predictable NIBS outcomes, several links between stimulation intensity and outcome variables remain under-researched, limiting the full potential of this approach. This includes rigorous and detailed assessments of behavioral and clinical relevance. We discuss these issues in detail, including the uncertainties associated with e-field computations and the challenges associated with transitioning from single-pulse to repetitive TMS thresholds. Addressing these knowledge gaps will help unlock the currently unexploited potential of TMS—and potentially other NIBS approaches—for the study of human cognition and the treatment of neurological and psychiatric disorders. Box 1 provides an overview of common implicit assumptions when using MT-based dosing outside the motor cortex.

## BOX 1

**Common assumptions of TMS dosing based on motor cortex excitability for non-motor areas**

MT- and e-field based TMS dosing individualize the stimulation strength based on motor-cortex excitability. When targeting non-motor regions, such as higher association cortices, with any of these dosing strategies several—usually implicit—assumptions are made about the mechanisms that underlie stimulation effects.

1. **Skin-cortex distance**: MT-based dosing assumes similar skin-cortex distances for the motor hotspot in the primary motor cortex (M1) and the stimulation target. Only for similar skin-cortex distances, cortical stimulation exposures are comparable across targets.
2. **Cortical stimulation thresholds**: All dosing strategies assume similar stimulation thresholds across the cortex, that is, the neuronal tissue at M1 and the stimulation target are assumed to have the same activation functions towards TMS pulses.
3. **Stimulation protocol:** Numerous studies use single-pulse TMS to quantify the motor cortex excitability but apply repetitive TMS (rTMS) to non-motor target areas, for example to modulate cognitive functions. Currently, all dosing strategies assume one global TMS threshold, independent of the temporal dynamics of the stimulation pattern (e.g., single-pulse TMS vs. rTMS).
4. **Outcome measure**: Many studies use similar cortical stimulation intensities (e.g. 100% of the resting motor threshold) to define motor cortex excitability (quantified via motor evoked potentials) and modulate behavioral responses (usually quantified as changes in response speed or accuracy) in cognitive experiments. Here, the same cortical stimulation threshold is assumed across different functional domains and outcome metrics. Similar rationales apply when using dosing strategies based on other cortical excitability estimates, such as the phosphene threshold in the visual cortex.

## Advantages of e-field-based dosing over motor threshold-based dosing

The motor threshold (MT) concept, which originated in the early days of TMS, was intended to standardize TMS effects across individuals and prevent adverse effects from overstimulation by individualizing the stimulation strength (Wassermann, 1998). The MT quantifies the *stimulator* device output (in % MSO) to yield muscle twitches (motor evoked potentials; MEPs) when stimulating the primary motor region (M1) with single TMS pulses. Despite its motor-centric definition, MT-based dosing is also commonly used for targets outside the motor cortex (Turi et al., 2021). A rationale for this generalization is the lack of a direct output measure to quantify the excitability of most non-motor regions. Critically, however, the *stimulator* output can only provide a rough estimate of the cortical *stimulation* exposure, which drives the cortical stimulation effects and is determined by the e-field magnitude and orientation at the cortex (Numssen et al., 2021). This poses various problems for MT-based dosing, most prominently for targets with a different skin-cortex distance than M1 (Koponen et al., 2020) or other macroscopic differences, such as different cortical thickness, yielding different cortical stimulation exposures for the same stimulator intensity. Likewise, the motor threshold does not correlate with the phosphene threshold (a metric for visual cortex excitability), when quantified via the *stimulator* intensity (Stewart et al., 2001; Boroojerdi et al., 2002). This illustrates that stimulation intensities for non-motor areas based on the motor threshold are somewhat arbitrary. E-field-based dosing approaches aim to solve this issue by normalizing the cortical *stimulation* strength across cortical areas. This can be achieved by computing the induced e-fields for both M1 and the actual target, and then determining the *stimulator* intensity required to produce the same cortical *stimulation* strength at the target area as in M1 at the motor threshold (see Caulfield et al., 2021a for a similar e-field dosing implementation):

*Intensity scaling factor = e-field at motor hotspot (in V/m) / e-field at target (in V/m)*
*Stimulator intensity (in % MSO) = MT (in % MSO) * Intensity scaling factor*

That is, the e-field strength in M1 at MT is used as a proxy for the individual *cortical stimulation threshold*—the cortical stimulation strength required to modulate neuronal processing with TMS.

To evaluate the effectiveness and feasibility of e-field based dosing across various cortical targets, we utilized this approach in 18 healthy participants (10 female; mean age = 30.5 years, SD = 5.88) to target different sensory-motor regions as well as higher-level association areas in the left hemisphere (Kuhnke & Numssen et al., 2023). Sensory-motor regions included M1, somatomotor and auditory cortices, while higher-level association regions included the inferior parietal lobe (IPL) and dorsolateral prefrontal cortex (DLPFC), which are common targets in TMS studies of cognition and for clinical applications (e.g. Beynel et al., 2019; Cash et al., 2021). This approach was ideal to elucidate potential advantages and drawbacks of e-field based dosing in comparison to MT-based dosing (e.g., Kuhnke et al., 2020) and other dosing strategies (Stokes et al., 2005).

Standard MT-based dosing applies the same *stimulator* intensity (in % MSO) at all (motor and non-motor) targets. Here, we used 100% resting MT intensity to allow for straightforward comparisons between targets, instead of applying a fraction of MT as is frequently done (e.g. 90% or 110% MT; Beynel et al, 2019; Kuhnke et al., 2017). As a second approach, we used a method proposed by Stokes et al. (2005) which aims to correct for cortical depth differences of motor- and non-motor-targets without assessing on the induced e-field. In Stokes’ approach, MT is adjusted by 3% MSO for every millimeter difference in skin-cortex distance between M1 and the target.

Like MT-based dosing and the Stokes approach, e-field based dosing depends on the correct localization of the hand muscle representation to accurately extract the cortical stimulation threshold. To this end, we employed a state-of-the-art TMS motor mapping protocol (Weise & Numssen et al., 2022b) to precisely locate the participant-specific first dorsal interosseous muscle representation in the primary motor cortex (M1) (see Supplemental Materials for details). Subsequently, we computed the optimal coil position for the motor hotspot and experimentally measured the resting MT with this coil position (i.e., the stimulator intensity required to elicit 5 out of 10 MEPs of size ≥ 50 µV). This allowed us to precisely quantify the cortical stimulation threshold (defined as resting MT) in terms of the cortical electric field strength in V/m at the motor hotspot instead of the hardware-dependent (Wang et al., 2023) stimulator intensity %MSO. Subsequently, we computed optimal coil positions for each of the four other targets (auditory, somatomotor, IPL, DLPFC). These non-motor targets were defined based on functional MRI localizers and meta-analyses, following standard procedures in current TMS studies on cognition. Please see *Remaining challenges – Target definitions* for potential future pathways to improve target selection and target delineation. Finally, we calculated the stimulator intensities required to elicit the same e-field strength in each target as in M1 at resting MT *–* the cortical stimulation threshold.

## Dosing comparison: Individual level

Figure 1 shows the comparison of the three dosing strategies for all targets in a representative individual participant. For M1 stimulation, the three dosing strategies do not differ as this stimulation target provides the common ground for the MT-based and e-field based approaches (Figure 1b & c, left column, Key Figure). M1 is effectively stimulated at 100% MT, leading to a hotspot of high cortical stimulation strength in the precentral gyrus. With MT-based dosing, the same stimulator intensity is simply applied to the other targets (e.g., 100% MT = 39% MSO for our example participant). While this approach seems to work reasonably well for the auditory cortex in this individual, the somatomotor cortex, IPL and DLPFC seem to be understimulated relative to M1 (Figure 1b, top row).

**Figure 1.**
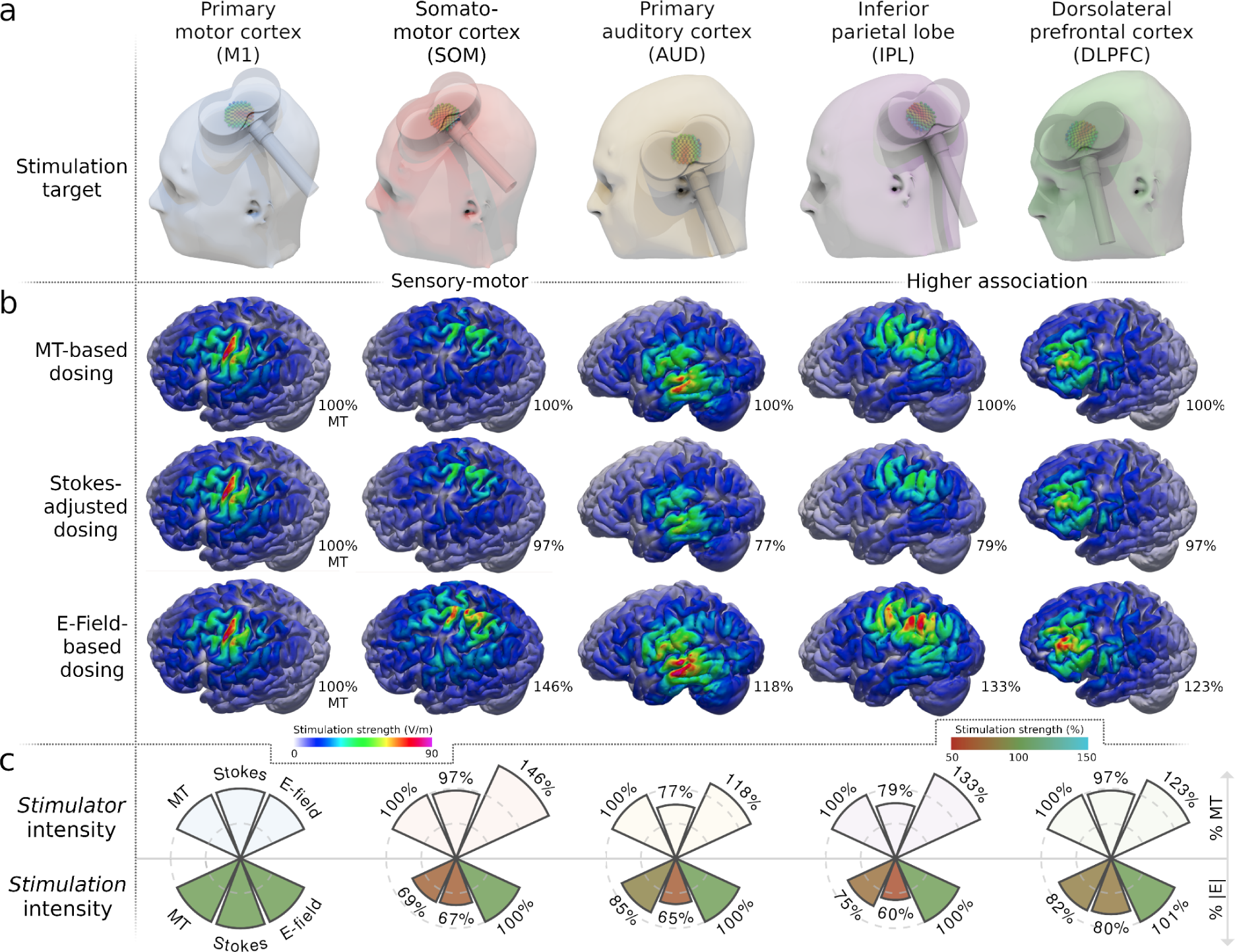
E-field dosing outperforms other dosing strategies on the participant level. (a) All coil placements were selected to maximize the cortical target stimulation. (b) Dosing based on the motor threshold (MT) alone (upper row) applies the same *stimulator* intensity across different cortical target regions (columns), yielding highly variable cortical stimulation strengths (quantified in volts per meter; V/m). The “Stokes” method (middle row) linearly adjusts the *stimulator* intensity for coil-to-target distance, but still results in a suboptimal match of cortical *stimulation* across targets. E-field based dosing (bottom row) yields the same cortical *stimulation* strength for all targets. Color: |E|. Percentages: % of MT stimulator intensity. All e-fields are visualized on the gray matter surface for one representative participant. (c) The relationship between *stimulator* intensity (upper row) and cortical *stimulation* exposure (bottom row) differs strongly across cortical targets. The *stimulation* exposures were extracted at the cortical targets and related to M1 exposure at MT intensity (‘100%’).

The Stokes approach yields lower stimulator intensities in our example participant for the non-motor targets (e.g., auditory: 77% MT; IPL: 79% MT) as these were located closer to the scalp than M1 (Figure 1b, middle row). Crucially, however, the Stokes approach also leads to understimulation of the non-motor areas in this participant (Figure 1b, middle row). The cortical stimulation exposure does not seem to be better matched between the non-motor targets and M1 than for MT-based dosing. Importantly, e-field based dosing leads to substantially different *stimulator* intensities for the different targets in our example participant (Figure 1b, bottom row). For instance, the somatomotor cortex needs to be stimulated at 146% MT (i.e., 57% MSO) to reach the cortical stimulation threshold for this participant.

Finally, e-field based dosing precisely matches the effective cortical stimulation for the different targets (Figure 1b, bottom row). For each target, e-field based dosing yields a hotspot of high cortical stimulation strength. Comparisons of the *stimulator* intensity and the cortical *stimulation* exposure identify strong differences between cortical regions (Figure 1c).

In conclusion, MT- and Stokes-based dosing fail to match the cortical stimulation intensity between different cortical targets within the same individual. In contrast, e-field dosing matches the cortical stimulation intensities for all the different targets (somatomotor, auditory, IPL, DLPFC) within our example participant to their individual cortical stimulation threshold (i.e., the e-field strength in M1 at rMT). Only e-field based dosing allows to equalize the stimulation exposure across different cortical areas within the same individual.

## Dosing comparison: Group level

Results on the group level (obtained from n = 18 healthy volunteers) parallel the central findings from the representative single-person case in Figure 1. MT-based and Stokes-adjusted dosing lead to strong variability of cortical stimulation strengths, both within individuals across targets and between individuals for the same target (Figure 2a). In contrast, e-field based dosing matches the cortical stimulation strength for each target to the respective participant’s cortical stimulation threshold—the individual stimulation strength in M1 at MT (Figure 2b)—minimizing within-participant variability. Minor deviations of the obtained e-field between targets within the same participant are caused by limitations of the stimulator device, whose intensity can only be changed in steps of 1% MSO. E-field dosing not only minimizes the within-participant variability but also reduces the between-participant variability, so that each cortical target receives the same stimulation strength on average (Figure 2a right).

**Figure 2.**
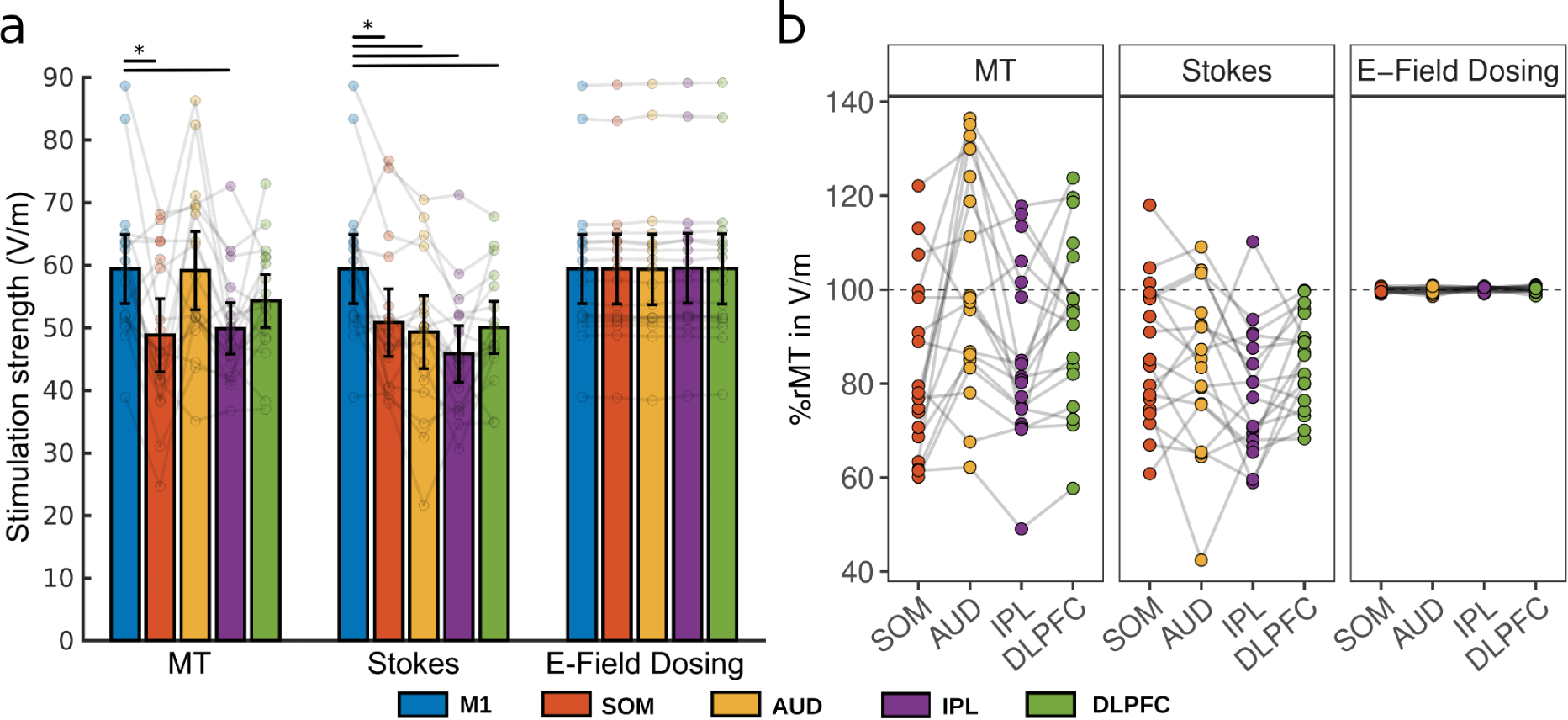
E-field based dosing outperforms other dosing strategies on the group level. (a) The cortical stimulation strength (in V/m) is highly variable for MT-based dosing (left) and Stokes-adjusted dosing (middle), both within and between the 18 participants across the different cortical targets. In contrast, e-field based dosing (right) yields the same stimulation strength for all targets within each participant, which also minimizes the between-participant variability. The cortical stimulation threshold centers around 60 V/m. (b) MT-based dosing fails to replicate cortical stimulation thresholds (y-axis) at other cortical targets and Stokes-adjusted dosing only slightly improves this relationship. E-field based dosing matches the cortical stimulation exposure at all cortical targets to the cortical stimulation threshold measured at M1. E-field magnitudes (|E|) were extracted and averaged within spherical ROIs (r = 5 mm) from gray matter volume only. Connected dots show individual participant data; error bars represent the 95% confidence interval. Black bars indicate significant differences (p < 0.05; Bonferroni-Holm corrected for multiple comparisons).

The average cortical stimulation threshold across participants was 59.5 V/m (SE = 2.8). For 15 out of 18 participants, the identified cortical threshold lies between 50 and 70 V/m, with one participant below and two participants above this range. Interestingly, cortical stimulation thresholds and model fits (between e-field and MEP magnitudes) are negatively correlated (r = -0.564, p = 0.019; see Figure S1). That is, participants with a better model fit have lower estimated cortical thresholds. Distinct differences in model fits might point to inaccuracies in the modeling pipeline or suboptimal data sampling. With several parameters of the modeling pipeline currently being subject to refinement, such as improved tissue segmentations, better group-based and individual estimates of tissue conductivities, and the impact of gray matter density on submillimeter e-field warps, the overall simulation accuracy will potentially be further improved in the future (see *Remaining challenges of e-field-based dosing - Simulation accuracy* section).

In conclusion, e-field based dosing better matches the cortical stimulation strength across the cortex than MT-based dosing approaches—both within and across individuals. Whereas e-field dosing matches the induced e-fields by design, MT and Stokes-based dosing fail to match the induced e-fields between different targets and individuals. Therefore, e-field based dosing may increase the stimulation efficacy and reduce both the within- and between-person variability of TMS effects. Overall, *a priori* e-field simulations promise to substantially improve TMS studies with non-motor target areas.

## Remaining challenges of e-field-based dosing

Despite significant improvements of e-field based dosing from conceptual and methodological perspectives, several challenges in the complex interplay between physiological, methodological, and cognitive parameters (Figure 3) remain to be addressed (Box “Outstanding Questions”). This includes assessment to test the behavioral and clinical relevance in experimental and clinical studies. Importantly, these crucial parameters, as detailed below, should not be considered in isolation but should instead be seen as a set of factors that together define the TMS-induced effects and may interact with each other.

**Figure 3.**
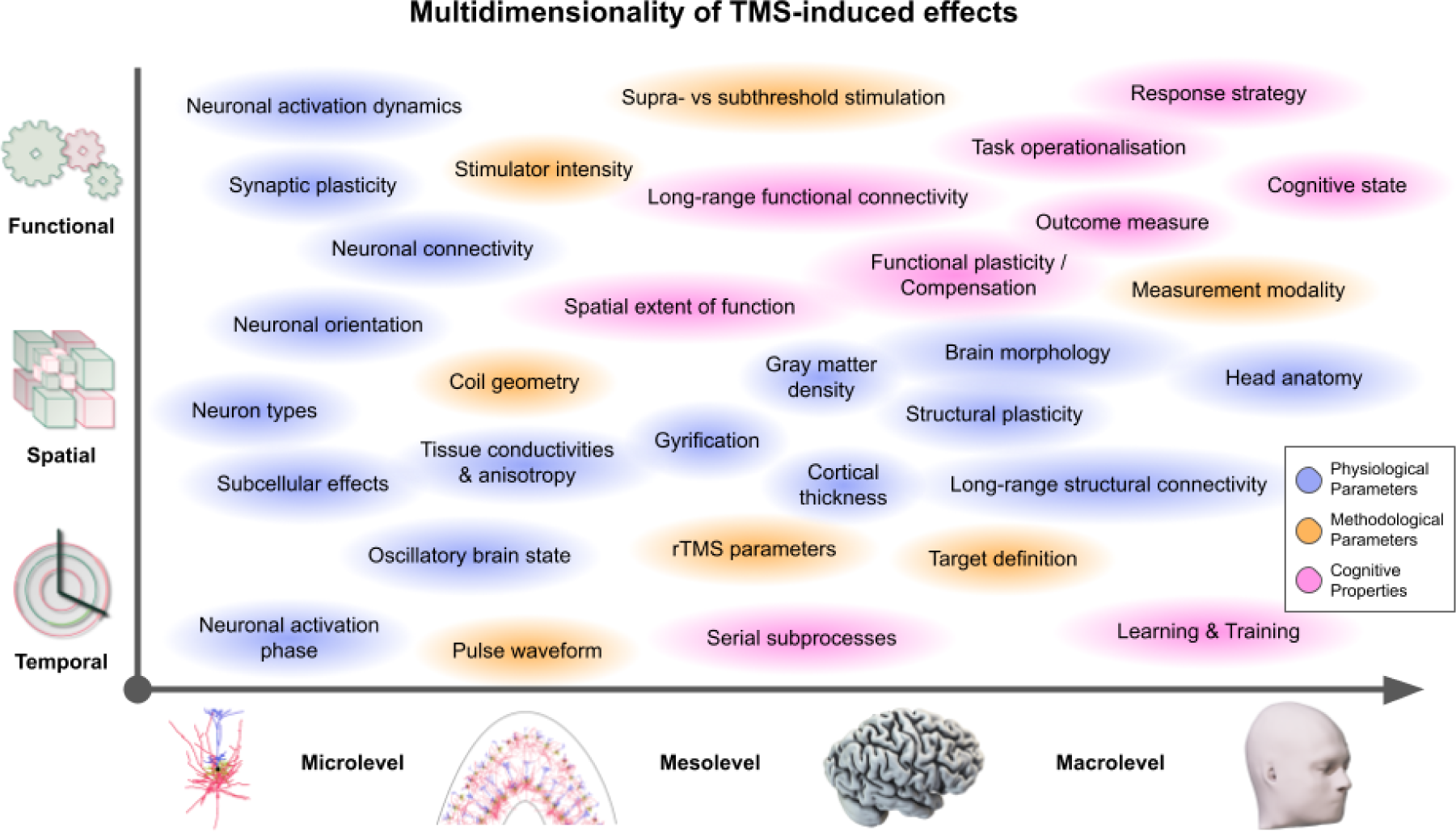
TMS-induced effects depend on a multidimensional set of factors. The outcome of a TMS study (or therapeutical intervention) is the sum of various factors, spanning **physiological parameters** (blue) of the individual (such as gyrification patterns and electric tissue properties), **methodological parameters** (orange; such as pulse waveform and target definition), and **cognitive properties** (magenta; such as response strategies and cognitive brain state). These are defined in the **temporal domain** (*microlevel*: different neuronal activation phases might yield different neuronal responses to a TMS pulse; *mesolevel*: stimulating during different serial subprocesses of a function; *macrolevel*: different levels of training render different cortical target regions effectively), the **spatial domain** (*microlevel*: neuronal orientation towards the induced e-field; *mesolevel*: the TMS coil geometry and gyrification patterns define the induced e-field; *macrolevel*: long-range white matter fiber tracts allow for distal TMS effects), or the **functional domain** (*microlev*el: single-cell activation mechanisms; *mesolevel*: supra-vs-subthreshold stimulation; *macrolevel*: outcome measure, such as response speed vs accuracy).

### Simulation accuracy

The development of easy-to-use toolboxes to compute the induced e-fields for individual anatomies and coil placements is still in its infancy and the field has not yet settled on common grounds. For example, the number of relevant tissue types that need to be assessed during e-field computation is still debated (Weise et al., 2022a). In addition, although considerable variations of relevant physiological properties such as electric conductivities across individuals and across the lifespan have been reported (Wagner et al., 2004; Antonakakis et al., 2020), these variations are currently not included in field modeling toolboxes. Approaches to estimate person-specific tissue conductivities *in vivo* are under development (e.g., magnetic resonance current density imaging; MRCDI; magnetic resonance electrical impedance tomography; MREIT; Göksu et al., 2018; Yazdanian et al., 2020; Eroğlu et al., 2021). Importantly, these variance sources do not necessarily impede within-participant comparisons and e-field dosing based on individualized M1 stimulation thresholds as these inaccuracies are constant within individuals and toolboxes. Instead, these limitations affect across-toolbox comparisons and considerations about mesoscopic stimulation mechanistics, which rely on physically correct e-field computations including precise information about the pulse shape. Due to these inaccuracies, the generalizability of individual cortical field thresholds, such as the ∼60 V/m estimate presented above, remains to be tested.

### Target definitions

E-field based dosing crucially depends on the correct definition of the cortical M1 region-of-interest (ROI) since the effective e-field that yields a neuronal effect is measured at this location. Specifically, the ROI location, parameters (ROI shape, size, etc.), and summary statistics (e.g. 99% percentile, maximum, etc.) influence this quantity and the brain stimulation community has yet to settle on common values for these parameters (see Van Hoornweder et al., 2023 for an overview on the different ROI definitions and extraction approaches used in TMS studies). Likewise, the “real” cortical target needs to be defined accurately, again, both with respect to its participant-specific location and spatial extent. However, in TMS studies of cognition, the cortical targets are often defined based on group-level fMRI or even meta-analyses (Beynel et al., 2019), ignoring individual differences, and thus, yielding suboptimal e-field calibrations. Besides participant-specific functional MRI localizers for TMS studies on cognition (Sack et al., 2008), promising strategies have been proposed that leverage individual functional connectivity and resting-state network mappings to define cortical targets (Lueckel et al., 2023; Lynch et al., 2022; Balderston et al., 2022). These approaches might also open the door towards defining off-targets, i.e. regions that should explicitly not be stimulated. Aside from a spatial target definition, the temporal target (Figure 3, bottom; Romei et al., 2016) also needs to be defined accurately to effectively modulate cortical processing of a specific function, potentially ranging from milliseconds (oscillatory states; Bergmann et al., 2012; Siebner et al., 2022) to seconds (sequential subprocesses of a function; Sack et al., 2005) to even longer time periods (adaptations due to learning and long-term plasticity or compensation; Bergmann & Hartwigsen, 2021).

### Different TMS protocols

Besides issues regarding the spatial distribution of the induced e-field, it is worth noting that while the motor threshold is typically assessed using single-pulse TMS, repetitive TMS (rTMS) is commonly employed in non-motor studies. For example, TMS for major depression therapy utilizes various different protocols, including 1 Hz to 20 Hz rTMS and several variations of theta-burst stimulation (Chen et al. 2013; Teng et al., 2017; Voigt et al., 2021). However, the mesoscopic effects of different rTMS protocols and, thus, their relation to MT thresholds based on single-pulse TMS are currently unknown. To gain a deeper understanding of this issue, modeling approaches will potentially provide valuable insights (see for example NeMo-TMS toolbox; Shirinpour et al., 2021). Currently, *cortical stimulation thresholds* are derived from a variety of experimental and modeling approaches, resulting in high variability. For example, experimental approaches identified thresholds from 35 V/m to 60 V/m when quantified via EEG (double pulses, Rosanova et al., 2009; individual alpha frequency, Zmeykina et al., 2020) and from 60 to above 100 V/m when measured with EMG (single pulses, Numssen et al., 2021; Kuhnke & Numssen et al., 2023). In contrast, modeling approaches and in vitro experiments identified thresholds between 120 V/m and 300 V/m (Turi et al., 2022; Shirinpour et al., 2021; Aberra et al., 2020, Weise et al., 2023, Aberra et al., 2023). It is important to note that modeling approaches currently use simplified setups, such as isolated neurons instead of interconnected neuronal networks and other cell types, and that neuronal models originate from rodent samples (Weise et al., 2023). Different pulse waveforms, such as mono-vs. biphasic (Wendt et al., 2023) and steeper vs. longer pulses (Sommer et al., 2023), add to the different neuronal activation dynamics (Peterchev et al., 2013; Koponen et al., 2018; Zeng et al., 2022). Further research in this area is needed to elucidate the relationships between cortical thresholds for single-pulse TMS and rTMS, particularly in non-motor regions.

### Different cortical targets

It is important to note that—besides the implicitly assumed similarity between single-pulse TMS and rTMS thresholds (see above)—also implicit assumptions of similar cortical responses for different cortical regions are made in (non-motor) TMS studies. For example, the cortical tissue of M1 and IPL differs substantially on the mesoscopic level, with M1 having large amounts of giant Betz cells in layer V and no layer IV, and the IPL showing the opposite pattern (Caspers et al., 2013). Despite this variability of physiological properties, cortical thresholds are currently assumed to be similar across different cortical regions. Most critically, recent modeling approaches have shown that different physiological properties (e.g. Dubbioso et al., 2021) and neuronal types, based on their morphology and orientation within the cortical tissue, show different coupling behavior to the induced electric field from TMS (Aberra et al., 2018, 2020; Weise et al., 2023). It should be noted that also macroscopic differences affect the induced e-field, including differences in cortical thickness, that require careful e-field quantification settings (e.g. surface vs. volume extraction). This questions the assumptions of similarity between cortical regions with respect to transcranial stimulation.

### Different functional domains

Finally, in TMS studies of cognition, the existence of a single cortical excitation threshold across different functional domains is assumed implicitly. The motor threshold measures the minimum cortical excitation (with single-pulse TMS of the primary motor region) necessary to elicit MEPs just above the EMG noise floor. However, it remains unknown if the same cortical excitation threshold can be applied to effectively modulate other (cognitive) functions, such as attentional reorienting (Jing et al., 2023), sentence processing (Meyer et al., 2018; Kuhnke et al., 2017; van der Burght, 2022) or conceptual-semantic processing (Kuhnke et al., 2020, 2023). As pre-activation of the motor cortex drastically lowers the cortical threshold to evoke MEPs (Rossi et al., 2009; see Holmes et al., 2023 for unintended side-effects), different (cognitive) brain states are also likely to affect stimulation effects within higher association cortices (Silvanto et al., 2008; Feurra et al., 2013; Krause et al., 2022; see Hartwigsen & Silvanto, 2022 for discussion). So far, relationships between functional domains and cortical excitation thresholds remain unknown.

## Concluding Remarks

Although e-field based dosing is no magic bullet for all challenges associated with NIBS studies, it has the potential to substantially decrease across- and within-person variance of cortical modulation in non-motor studies. By providing a more biologically plausible dosing metric, this approach can potentially play a crucial role in improving the personalization of TMS and tES treatments in clinical settings, as well as increasing the effect sizes of NIBS studies at the group level. Crucially, these assumptions have yet to be tested in experimental and clinical studies. As such, e-field based dosing represents a promising avenue for future research in the field of NIBS.

## Data and Code Availability

Electric field values from the regions of interest and analysis scripts are made public in an online repository.

## Author Contributions

**Ole Numssen:** Conceptualization, Data curation, Formal analysis, Investigation, Methodology, Software, Validation, Visualization, Writing – original draft, Writing – review & editing; **Philipp Kuhnke:** Conceptualization, Data curation, Formal analysis, Methodology, Validation, Visualization, Writing – original draft, Writing – review & editing; **Konstantin Weise:** Conceptualization, Methodology, Validation, Writing – original draft, Writing – review & editing; **Gesa Hartwigsen:** Conceptualization, Project administration, Resources, Supervision, Writing – original draft, Writing – review & editing

## References

Aberra, A. S., Peterchev, A. V., & Grill, W. M. (2018). Biophysically realistic neuron models for simulation of cortical stimulation. J. Neural Eng., 15(6). DOI: 10.1088/1741-2552/aadbb1

Aberra, A. S., Wang, B., Grill, W. M., & Peterchev, A. V. (2020). Simulation of transcranial magnetic stimulation in head model with morphologically-realistic cortical neurons. Brain Stimul., 13(1), 175–189. DOI: 10.1016/j.brs.2019.10.002

Aberra, A. S., Lopez, A., Grill, W. M., & Peterchev, A. V. (2023). Rapid estimation of cortical neuron activation thresholds by transcranial magnetic stimulation using convolutional neural networks. NeuroImage, 275, 120184.

Antonakakis, M., Schrader, S., Aydin, Ü., Khan, A., Gross, J., Zervakis, M., Rampp, S., & Wolters, C. H. (2020). Inter-subject variability of skull conductivity and thickness in calibrated realistic head models. NeuroImage, 223, 117353. DOI: 10.1016/j.neuroimage.2020.117353

Balderston, N. L., Beer, J. C., Seok, D., Makhoul, W., Deng, Z. D., Girelli, T., … & Sheline, Y. I. (2022). Proof of concept study to develop a novel connectivity-based electric-field modelling approach for individualized targeting of transcranial magnetic stimulation treatment. Neuropsychopharmacology, 47(2), 588–598. DOI: 10.1038/s41386-021-01110-6

Bergmann, T. O., Mölle, M., Schmidt, M. A., Lindner, C., Marshall, L., Born, J., & Siebner, H. R. (2012). EEG-guided transcranial magnetic stimulation reveals rapid shifts in motor cortical excitability during the human sleep slow oscillation. J. Neurosci., 32(1), 243–253. DOI: 10.1523/JNEUROSCI.4792-11.2012

Bergmann, T. O., & Hartwigsen, G. (2021). Inferring causality from noninvasive brain stimulation in cognitive neuroscience. J. Cogn. Neurosci., 33(2), 195–225. DOI: 10.1162/jocn_a_01591

Beynel, L., Appelbaum, L. G., Luber, B., Crowell, C. A., Hilbig, S. A., Lim, W., Nguyen, D., Chrapliwy, N. A., Davis, S. W., Cabeza, R., Lisanby, S. H., & Deng, Z. D. (2019). Effects of online repetitive transcranial magnetic stimulation (rTMS) on cognitive processing: A meta-analysis and recommendations for future studies. Neurosci. Biobehav. Rev., 107, 47–58. DOI: 10.1016/j.neubiorev.2019.08.018

Boroojerdi, B., Meister, I. G., Foltys, H., Sparing, R., Cohen, L. G., & Töpper, R. (2002). Visual and motor cortex excitability: a transcranial magnetic stimulation study. Clin. Neurophysiol., 113(9), 1501–1504. DOI: 10.1016/s1388-2457(02)00198-0

Burt, T., Lisanby, S. H., & Sackeim, H. A. (2002). Neuropsychiatric applications of transcranial magnetic stimulation: a meta analysis. International Journal of Neuropsychopharmacology, 5(1), 73–103.

Caulfield, K. A., Li, X., & George, M. S. (2021a). Four electric field modeling methods of Dosing Prefrontal Transcranial Magnetic Stimulation (TMS): Introducing APEX MT dosimetry. Brain Stimul., 14(4), 1032–1034. DOI: 10.1016/j.brs.2021.06.012

Caulfield, K. A., Li, X., & George, M. S. (2021b). A reexamination of motor and prefrontal TMS in tobacco use disorder: Time for personalized dosing based on electric field modeling?. Clinical Neurophysiology, 132(9), 2199–2207.

Caulfield, K. A., & Brown, J. C. (2022). The problem and potential of TMS’ infinite parameter space: a targeted review and road map forward. Frontiers in Psychiatry, 13.

Cash, R. F., Weigand, A., Zalesky, A., Siddiqi, S. H., Downar, J., Fitzgerald, P. B., & Fox, M. D. (2021). Using brain imaging to improve spatial targeting of transcranial magnetic stimulation for depression. Biol. Psychiatry, 90(10), 689–700. DOI: 10.1016/j.biopsych.2020.05.033

Caspers, S., Schleicher, A., Bacha-Trams, M., Palomero-Gallagher, N., Amunts, K., & Zilles, K. (2013). Organization of the human inferior parietal lobule based on receptor architectonics. Cereb. Cortex, 23(3), 615–628.DOI: 10.1093/cercor/bhs048

Chen, J., Zhou, C., Wu, B., Wang, Y., Li, Q., Wei, Y., Yang, D., Mu, J., Zhu, D., Zou, D., & Xie, P. (2013). Left versus right repetitive transcranial magnetic stimulation in treating major depression: a meta-analysis of randomised controlled trials. Psychiatry Res., 210(3), 1260–1264. DOI: 10.1016/j.psychres.2013.09.007

Dannhauer, M., Huang, Z., Beynel, L., Wood, E., Bukhari-Parlakturk, N., & Peterchev, A. V. (2022). TAP: Targeting and analysis pipeline for optimization and verification of coil placement in transcranial magnetic stimulation. J. Neural Eng., 19(2). DOI: 10.1088/1741-2552/ac63a4

Dubbioso, R., Madsen, K. H., Thielscher, A., & Siebner, H. R. (2021). The myelin content of the human precentral hand knob reflects interindividual differences in manual motor control at the physiological and behavioral level. J. Neurosci., 41(14), 3163–3179. DOI: 10.1523/JNEUROSCI.0390-20.2021

Eroğlu, H. H., Puonti, O., Göksu, C., Gregersen, F., Siebner, H. R., Hanson, L. G., & Thielscher, A. (2021). On the reconstruction of magnetic resonance current density images of the human brain: Pitfalls and perspectives. NeuroImage, 243, 118517. DOI: 10.1016/j.neuroimage.2021.118517

Esmaeilpour, Z., Kronberg, G., Reato, D., Parra, L. C., & Bikson, M. (2021). Temporal interference stimulation targets deep brain regions by modulating neural oscillations. Brain Stimulation, 14(1), 55–65.

Feurra, M., Pasqualetti, P., Bianco, G., Santarnecchi, E., Rossi, A., & Rossi, S. (2013). State-dependent effects of transcranial oscillatory currents on the motor system: what you think matters. J. Neurosci., 33(44), 17483–17489. DOI: 10.1523/JNEUROSCI.1414-13.2013

Göksu, C., Hanson, L. G., Siebner, H. R., Ehses, P., Scheffler, K., & Thielscher, A. (2018). Human in-vivo brain magnetic resonance current density imaging (MRCDI). NeuroImage, 171, 26–39. DOI: 10.1016/j.neuroimage.2017.12.075

Hamada, M., Murase, N., Hasan, A., Balaratnam, M., & Rothwell, J. C. (2013). The role of interneuron networks in driving human motor cortical plasticity. Cereb. Cortex, 23(7), 1593–1605. DOI: 10.1093/cercor/bhs147

Hartwigsen, G., & Silvanto, J. (2022). Noninvasive brain stimulation: Multiple effects on cognition. The Neuroscientist, 10738584221113806. DOI:10.1177/10738584221113806

Hartwigsen, G., Bergmann, T. O., Herz, D. M., Angstmann, S., Karabanov, A., Raffin, E., Thielscher, A., & Siebner, H. R. (2015). Modeling the effects of noninvasive transcranial brain stimulation at the biophysical, network, and cognitive Level. Prog. Brain Res., 222, 261–287. DOI: 10.1016/bs.pbr.2015.06.014

Holmes, N. P., Di Chiaro, N. V., Crowe, E., & Reader, A. T.(2023). TMS over parietal and premotor cortex evokes MEPs from motor cortex. OSF preprint. DOI:0.17605/OSF.IO/2XYTM

Huang, Y., Datta, A., Bikson, M., & Parra, L. C. (2019). Realistic volumetric-approach to simulate transcranial electric stimulation—ROAST—a fully automated open-source pipeline. J. Neural Eng., 16(5), 056006. DOI: 10.1088/1741-2552/ab208d

Jing, Y., Numssen, O., Weise, K., Kalloch, B., Buchberger, L., Haueisen, J., Hartwigsen, G., & Knoesche, T. (2023). Modeling the Effects of Transcranial Magnetic Stimulation on Spatial Attention. bioRxiv. DOI: 10.1101/2023.01.11.523548

Kasten, F. H., Duecker, K., Maack, M. C., Meiser, A., & Herrmann, C. S. (2019). Integrating electric field modeling and neuroimaging to explain inter-individual variability of tACS effects. Nat. Commun., 10, DOI: 10.1038/s41467-019-13417-6

Koponen, L. M., Nieminen, J. O., Mutanen, T. P., & Ilmoniemi, R. J. (2018). Noninvasive extraction of microsecond-scale dynamics from human motor cortex. Hum. Brain Mapp., 39(6), 2405–2411. DOI: 10.1002/hbm.24010

Koponen, L. M., & Peterchev, A. V. (2020). Transcranial magnetic stimulation: principles and applications. Neural Engineering, 245-270.

Krause, M. R., Vieira, P. G., Thivierge, J. P., & Pack, C. C. (2022). Brain stimulation competes with ongoing oscillations for control of spike timing in the primate brain. PLoS Biol., 20(5), e3001650. DOI: 10.1371/journal.pbio.3001650

Kuhnke, P., Beaupain, M.C., Cheung, V.K.M., Weise, K., Kiefer, M., & Hartwigsen, G. (2020). Left posterior inferior parietal cortex causally supports the retrieval of action knowledge. NeuroImage, 219, 117041. DOI: 10.1016/j.neuroimage.2020.117041

Kuhnke, P., Beaupain, M. C., Arola, J., Kiefer, M., & Hartwigsen, G. (2023). Meta-analytic evidence for a novel hierarchical model of conceptual processing. Neuroscience & Biobehavioral Reviews, 144, 104994. 10.1016/j.neubiorev.2022.104994

Kuhnke, P., Numssen, O., Voeller, J., Weise, K., & Hartwigsen, G. (2023). P-87 Dosage optimization for transcranial magnetic stimulation based on cortical field thresholds. Clin. Neurophysiol., 148, e48. DOI: 10.1016/j.clinph.2023.02.104

Kuhnke, P., Meyer, L., Friederici, A.D., & Hartwigsen, G. (2017). Left posterior inferior frontal gyrus is causally involved in reordering during sentence processing. NeuroImage, 148, 254–263. DOI: 10.1016/j.neuroimage.2017.01.013

Laakso I, Mikkonen M, Koyama S, Hirata A, & Tanaka S (2019). Can electric fields explain inter-individual variability in transcranial direct current stimulation of the motor cortex? Sci. Rep., 9, 626. DOI: 10.1038/s41598-018-37226-x

Lee, J. C., Corlier, J., Wilson, A. C., Tadayonnejad, R., Marder, K. G., Ngo, D., Krantz, D. E., Wilke, S. A., Levitt, J. G., Ginder, N. D., & Leuchter, A. F. (2021). Subthreshold stimulation intensity is associated with greater clinical efficacy of intermittent theta-burst stimulation priming for Major Depressive Disorder. Brain Stimul., 14(4), 1015–1021. DOI: 10.1016/j.brs.2021.06.008

Lueckel, M., Radetz, A., Hassan, U., Yuen, K., Mueller-Dahlhaus, F., Kalisch, R., & Bergmann, T. O. (2023). Functional connectivity-and E-field-optimized TMS targeting: method development and concurrent TMS-fMRI validation. Brain Stimulation: Basic, Translational, and Clinical Research in Neuromodulation, 16(1), 189. DOI: 10.1016/j.brs.2023.01.223

Lynch, C. J., Elbau, I. G., Ng, T. H., Wolk, D., Zhu, S., Ayaz, A., … & Liston, C. (2022). Automated optimization of TMS coil placement for personalized functional network engagement. Neuron, 110(20), 3263–3277. DOI: 10.1016/j.neuron.2022.08.012

Meyer, L., Elsner, A., Turker, S., Kuhnke, P., & Hartwigsen, G. (2018). Perturbation of left posterior prefrontal cortex modulates top-down processing in sentence comprehension. NeuroImage, 181(July), 598–604. 10.1016/j.neuroimage.2018.07.059

Numssen, O., van der Burght, C. L., & Hartwigsen, G. (2023). Revisiting the focality of non-invasive brain stimulation – Implications for studies of human cognition. Neurosci. Biobehav. Rev., 149, 105154. DOI: 10.1016/j.neubiorev.2023.105154

Numssen, O., Zier, A. L., Thielscher, A., Hartwigsen, G., Knösche, T. R., & Weise, K. (2021). Efficient high-resolution TMS mapping of the human motor cortex by nonlinear regression. NeuroImage, 245, 118654. DOI: 10.1016/j.neuroimage.2021.118654

O’Reardon, J. P., Solvason, H. B., Janicak, P. G., Sampson, S., Isenberg, K. E., Nahas, Z., … & Sackeim, H. A. (2010). Reply regarding “efficacy and safety of transcranial magnetic stimulation in the acute treatment of major depression: a multisite randomized controlled trial”. Biological psychiatry, 67(2), e15–e17.

Pascual-Leone, A. (2000). Transcranial magnetic stimulation in cognitive neuroscience – virtual lesion, chronometry, and functional connectivity. Curr. Opin. Neurobiol., 10(2), 232–237. DOI: 10.1016/S0959-4388(00)00081-7

Peterchev, A. V., Goetz, S. M., Westin, G. G., Luber, B., & Lisanby, S. H. (2013). Pulse width dependence of motor threshold and input–output curve characterized with controllable pulse parameter transcranial magnetic stimulation. Clin. Neurophysiol., 124(7), 1364–1372. DOI: 10.1016/j.clinph.2013.01.011

Peterchev, A. V., Wagner, T. A., Miranda, P. C., Nitsche, M. A., Paulus, W., Lisanby, S. H., … & Bikson, M. (2012). Fundamentals of transcranial electric and magnetic stimulation dose: definition, selection, and reporting practices. Brain stimulation, 5(4), 435–453. DOI: 10.1016/j.brs.2011.10.001

Puonti, O., Van Leemput, K., Saturnino, G. B., Siebner, H. R., Madsen, K. H., & Thielscher, A. (2020). Accurate and robust whole-head segmentation from magnetic resonance images for individualized head modeling. NeuroImage, 219, 117044. DOI: 10.1016/j.neuroimage.2020.117044

Romei, V., Thut, G., & Silvanto, J. (2016). Information-based approaches of noninvasive transcranial brain stimulation. Trends Neurosci., 39(11), 782–795. DOI: 10.1016/j.tins.2016.09.001

Rosanova, M., Casali, A., Bellina, V., Resta, F., Mariotti, M., & Massimini, M. (2009). Natural Frequencies of Human Corticothalamic Circuits. J. Neurosci., 29(24), 7679–7685. DOI: 10.1523/JNEUROSCI.0445-09.2009

Rossi, S., Hallett, M., Rossini, P. M., Pascual-Leone, A., & Safety of TMS Consensus Group (2009). Safety, ethical considerations, and application guidelines for the use of transcranial magnetic stimulation in clinical practice and research. Clin. Neurophysiol., 120(12), 2008–2039. DOI: 10.1016/j.clinph.2009.08.016

Rossini, P. M., Barker, A. T., Berardelli, A., Caramia, M. D., Caruso, G., Cracco, R. Q., Dimitrijević, M. R., Hallett, M., Katayama, Y., Lücking, C. H., Maertens de Noordhout, A. L., Marsden, C.D., Murray, N.M.F., Rothwell, J.C., Swash, M., Tomberg, C. (1994). Non-invasive electrical and magnetic stimulation of the brain, spinal cord and roots: basic principles and procedures for routine clinical application. Report of an IFCN committee. Electroencephalogr. Clin. Neurophysiol., 91(2), 79–92. DOI: 10.1016/0013-4694(94)90029-9

Sack, A. T., Camprodon, J. A., Pascual-Leone, A., & Goebel, R. (2005). The dynamics of interhemispheric compensatory processes in mental imagery. Science, 308(5722), 702–704. DOI: 10.1126/science.1107784

Sack, A. T., Kadosh, R. C., Schuhmann, T., Moerel, M., Walsh, V., & Goebel, R. (2009). Optimizing functional accuracy of TMS in cognitive studies: a comparison of methods. Journal of cognitive neuroscience, 21(2), 207–221.

Sandrini, M., Umiltà, C., & Rusconi, E. (2011). The use of transcranial magnetic stimulation in cognitive neuroscience: A new synthesis of methodological issues. Neurosci. Biobehav. Rev., 35, 516–536. DOI: 10.1016/j.neubiorev.2010.06.005

Sasaki, T., Kodama, S., Togashi, N., Shirota, Y., Sugiyama, Y., Tokushige, S., Inomata-Terada, S., Terao, Y., Ugawa, Y., Hamada, M. (2018). The intensity of continuous theta burst stimulation, but not the waveform used to elicit motor evoked potentials, influences its outcome in the human motor cortex. Brain Stimul., 11(2), 400–410. DOI: 10.1016/j.brs.2017.12.003

Saturnino, G. B., Puonti, O., Nielsen, J. D., Antonenko, D., Madsen, K. H., & Thielscher, A. (2019). SimNIBS 2.1: A Comprehensive Pipeline for Individualized Electric Field Modelling for Transcranial Brain Stimulation. In: Makarov, S., Horner, M., Noetscher, G. (Eds.), Brain and Human Body Modeling (3–25). Springer, Cham. DOI: 10.1007/978-3-030-21293-3_1

Shirinpour, S., Hananeia, N., Rosado, J., Tran, H., Galanis, C., Vlachos, A., Jedlicka, P., Queisser, G., & Opitz, A. (2021). Multi-scale modeling toolbox for single neuron and subcellular activity under Transcranial Magnetic Stimulation. Brain Stimul., 14(6), 1470–1482. DOI: 10.1016/j.brs.2021.09.004

Siebner, H. R., Funke, K., Aberra, A. S., Antal, A., Bestmann, S., Chen, R., Classen, J., Davare, M., Di Lazzaro, V., Fox, P. T., Hallett, M., Karabanov, A. N., Kesselheim, J., Beck, M. M., Koch, G., Liebetanz, D., Meunier, S., Miniussi, C., Paulus, W., Peterchev, A. V., … Ugawa, Y. (2022). Transcranial magnetic stimulation of the brain: What is stimulated? – A consensus and critical position paper. Clin. Neurophysiol., 140, 59–97. DOI: 10.1016/j.clinph.2022.04.022

Silvanto, J., Muggleton, N., & Walsh, V. (2008). State-dependency in brain stimulation studies of perception and cognition. Trends Cogn. Sci., 12(12), 447–454. DOI: 10.1016/j.tics.2008.09.004

Sommer, M., & Paulus, W. (2023). TMS waveform and current direction. The Oxford handbook of transcranial stimulation (2nd Edition). Oxford University Press, New York, 7–12. DOI: 10.1093/oxfordhb/9780198832256.013.5

Stewart, L. M., Walsh, V., & Rothwell, J. C. (2001). Motor and phosphene thresholds: a transcranial magnetic stimulation correlation study. Neuropsychologia, 39(4), 415–419. DOI: 10.1016/s0028-3932(00)00130-5

Stokes, M. G., Chambers, C. D., Gould, I. C., Henderson, T. R., Janko, N. E., Allen, N. B., & Mattingley, J. B. (2005). Simple Metric For Scaling Motor Threshold Based on Scalp-Cortex Distance: Application to Studies Using Transcranial Magnetic Stimulation. J. Neurophysiol., 94, 4520–4527. DOI: 10.1152/jn.00067.2005

Teng, S., Guo, Z., Peng, H., Xing, G., Chen, H., He, B., McClure, M. A., & Mu, Q. (2017). High-frequency repetitive transcranial magnetic stimulation over the left DLPFC for major depression: session-dependent efficacy: a meta-analysis. Eur. Psychiatry, 41(1), 75–84. DOI: 10.1016/j.eurpsy.2016.11.002

Thielscher, A., Antunes, A., & Saturnino, G. B. (2015). Field modeling for transcranial magnetic stimulation: A useful tool to understand the physiological effects of TMS? Annu. Int. Conf. IEEE Eng. Med. Biol. Soc., 2015, 222–225. DOI: 10.1109/embc.2015.7318340

Turi, Z., Lenz, M., Paulus, W., Mittner, M., & Vlachos, A. (2021). Selecting stimulation intensity in repetitive transcranial magnetic stimulation studies: A systematic review between 1991 and 2020. Eur. J. Neurosci., 53(10), 3404–3415. DOI: 10.1111/ejn.15195

Turi, Z., Hananeia, N., Shirinpour, S., Opitz, A., Jedlicka, P., & Vlachos, A. (2022). Dosing Transcranial Magnetic Stimulation of the Primary Motor and Dorsolateral Prefrontal Cortices With Multi-Scale Modeling. Frontiers in Neuroscience, 16(July), 1–11. DOI: 10.3389/fnins.2022.929814

van der Burght, C. L., Numssen, O., Schlaak, B., Goucha, T., & Hartwigsen, G. (2023). Differential contributions of inferior frontal gyrus subregions to sentence processing guided by intonation. Human Brain Mapping, 44(2), 585–598. 10.1002/hbm.26086

Van Hoornweder, S., Nuyts, M., Frieske, J., Verstraelen, S., Meesen, R. L. J., & Caulfield, K. A. (2023). A systematic review and large-scale tES and TMS electric field modeling study reveals how outcome measure selection alters results in a person- and montage- specific manner. BioRxiv. DOI: 10.1101/2023.02.22.529540

Voigt, J. D., Leuchter, A. F., & Carpenter, L. L. (2021). Theta burst stimulation for the acute treatment of major depressive disorder: A systematic review and meta-analysis. Transl. Psychiatry, 11(1), 330. DOI: 10.1038/s41398-021-01441-4

Wagner, T. A., Zahn, M., Grodzinsky, A. J., & Pascual-Leone, A. (2004). Three-dimensional head model simulation of transcranial magnetic stimulation. IEEE Trans. Biomed. Eng., 51(9), 1586–1598. DOI: 10.1109/TBME.2004.827925

Walsh, V., & Cowey, A. (2000). Transcranial magnetic stimulation and cognitive neuroscience. Nat. Rev. Neurosci., 1(1), 73–79. DOI: 10.1038/35036239

Wang, Y., Vora, I., Huynh, B. P., Picard-Fraser, M., Daneshzand, M., Nummenmaa, A., & Kimberley, T. J. (2023). Coils are not created equal: Effects on TMS thresholding. *Brain Stimulation: Basic*, Translational, and Clinical Research in Neuromodulation.

Wassermann, E. M. (1998). Risk and safety of repetitive transcranial magnetic stimulation: report and suggested guidelines from the International Workshop on the Safety of Repetitive Transcranial Magnetic Stimulation, June 5–7, 1996. Electroencephalogr. Clin. Neurophysiol., 108, 1–16. DOI: 10.1016/S0168-5597(97)00096-8

Weise, K., Numssen, O., Thielscher, A., Hartwigsen, G., & Knösche, T. R. (2020). A novel approach to localize cortical TMS effects. NeuroImage, 209, 116486. DOI: 10.1016/j.neuroimage.2019.116486

Weise, K., Wartman, W. A., Knösche, T. R., Nummenmaa, A. R., & Makarov, S. N. (2022a). The effect of meninges on the electric fields in TES and TMS. Numerical modeling with adaptive mesh refinement. Brain Stimul., 15(3), 654–663. DOI: 10.1016/j.brs.2022.04.009

Weise, K., Numssen, O., Kalloch, B., Zier, A. L., Thielscher, A., Haueisen, J., Hartwigsen, G., & Knösche, T. R. (2022b). Precise motor mapping with transcranial magnetic stimulation. Nat. protoc., 18(2), 293–318. DOI: 10.1038/s41596-022-00776-6

Weise, K., Worbs, T. H., Kalloch, B., Souza, V. H., Jaquier, A. T., Van Geit, W., … & Knösche, T. R. (2023). Directional Sensitivity of Cortical Neurons Towards TMS Induced Electric Fields. Imaging Neuroscience. DOI: 10.1162/imag_a_00036

Wendt, K., Sorkhabi, M. M., Stagg, C. J., Fleming, M. K., Denison, T., & O’Shea, J. (2023). The effect of pulse shape in theta-burst stimulation: Monophasic vs biphasic TMS. Brain Stimulation, 16(4), 1178–1185.

Wischnewski, M., Mantell, K. E., & Opitz, A. (2021). Identifying regions in prefrontal cortex related to working memory improvement: a novel meta-analytic method using electric field modeling. Neurosci. Biobehav. Rev., 130, 147–161. DOI: 10.1016/j.neubiorev.2021.08.017

Yazdanian, H., Saturnino, G. B., Thielscher, A., & Knudsen, K. (2020). Fast evaluation of the Biot-Savart integral using FFT for electrical conductivity imaging. J. Comput. Phys., 411, 109408. DOI: 10.1016/j.jcp.2020.109408

Zeng, Z., Koponen, L. M., Hamdan, R., Li, Z., Goetz, S. M., & Peterchev, A. V. (2022). Modular multilevel TMS device with wide output range and ultrabrief pulse capability for sound reduction. J. Neural Eng., 19(2), 026008. DOI: 10.1088/1741-2552/ac572c

Zmeykina, E., Mittner, M., Paulus, W., & Turi, Z. (2020). Weak rTMS-induced electric fields produce neural entrainment in humans. Sci. Rep., 10(1), 11994. DOI: 10.1038/s41598-020-68687-8

