## Supplemental Material for "Electric field based dosing for TMS"

* shared first authors

### Locating the motor hotspot with TMS

Like MT-based dosing and the Stokes approach, e-field based dosing depends on the correct localization of the hand muscle representation to accurately extract the cortical stimulation threshold. To this end, we employed a state-of-the-art TMS motor mapping protocol (Weise & Numssen et al., 2022) to precisely localize the participant-specific muscle representation in the primary motor cortex (M1). This protocol utilizes pyNIBS v0.3 (Numssen et al., 2021) and SimNIBS v4.0 (Saturnino et al., 2019; Thielscher et al., 2015): First, we constructed high-resolution head models based on individual T1-, T2-, and diffusion-weighted MR images (for details, see *Electric field modeling details* below). Second, we applied ~300 TMS pulses in a 2-3 cm radius around the presumptive motor hotpot, and recorded the coil positions/orientations and motor evoked potentials (MEPs) from the first dorsal interosseous muscle. Third, we simulated the TMS-induced e-field for each pulse using SimNIBS. Fourth, for each cortical element (in a region-of-interest of M1, premotor, and somatosensory cortices), we fit the local e-field magnitude onto the MEP magnitude and quantified the goodness-of-fit ($R^2$). Fifth, we identified the motor hotspot as the cortical element with the best fit (highest $R^2$) on the M1 gyral crown. Sixth, we computed the optimal coil position for the motor hotspot. To this end, we selected the coil position from a grid of 2-cm radius and 2-mm steps that maximized the e-field magnitude in a 5-mm sphere around the motor hotspot, restricted to gray matter ('position optimization'; see Numssen et al., 2021 for details). Finally, we experimentally measured the resting MT at this optimal coil position (i.e., the stimulator intensity required to elicit 5 out of 10 MEPs of size ≥ 50 μV). This allowed us to precisely quantify the cortical stimulation threshold (defined as resting MT) in V/m at the cortical motor hotspot.

### Electric field modeling details

For the electric field simulations with the finite element method (FEM, SimNIBS v4.0, Saturnino et al., 2019) we created participant-specific high-resolution head models based on individual T1-, T2-, and diffusion-weighted MR images (using the CHARM method; Puonti et al., 2020).

MR images were acquired using a 3T MRI scanner (Siemens, Erlangen, Germany) using the following sequences:

- **T1-weighted MPRAGE**: 176 slices in sagittal orientation; repetition time (TR): 2.3 s; echo time (TE): 2.98 ms; field of view: 256 mm; voxel size: 1 x 1 x 1 mm; no slice gap; flip angle: 9°; phase encoding direction: A/P
- **T2-weighted**: 192 slices in sagittal orientation; voxel size = 0.9 x 0.45 x 0.45 mm; flip angle = 120°; TR = 3.2 s; TE = 408 ms
- **diffusion-weighted**: 67 slices in axial orientation; matrix size = 128 x 128; voxel size = 1.7 x 1.7 x 1.7 mm; flip angle = 90°; TR = 7 s; TE = 80 ms, 67 diffusion directions, b-value 1000 s/mm3

T1- and T2-weighted images were used for segmenting the main tissues of the head: scalp, skull, grey matter (GM), white matter (WM), and cerebrospinal fluid (CSF). Diffusion-weighted images were employed to include anisotropic conductivity information. The electrical field was calculated for 1 A/μs, using default tissue conductivities values ($\sigma_{scalp}$ = 0.465 S/m, $\sigma_{skull}$ = 0.01 S/m, $\sigma_{GM}$ = 0.275 S/m, $\sigma_{WM}$ = 0.126 S/m, $\sigma_{CSF}$ = 1.654 S/m; Thielscher et al., 2011; Wagner et al., 2004).

#### Sample Information

**Table S1.** Sample statistics - Individual participants

| ID | Sex | Age | LQ | rMT | SOM Ratio | AUD Ratio | IPL Ratio | DLPFC Ratio | SOM % | AUD % | IPL % | DLPFC % |
|---|---|---|---|---|---|---|---|---|---|---|---|---|
| 01 | M | 37 | 100 | 40 | 1.350 | 0.730 | 0.850 | 0.840 | 54 | 29 | 34 | 33 |
| 02 | M | 40 | 100 | 42 | 1.630 | 1.610 | 2.040 | 1.730 | 68 | 68 | 86 | 73 |
| 03 | M | 38 | 75 | 49 | 1.280 | 0.840 | 1.020 | 1.080 | 63 | 41 | 50 | 53 |
| 04 | F | 33 | 80 | 35 | 1.580 | 0.750 | 0.880 | 1.020 | 55 | 26 | 31 | 36 |
| 05 | F | 37 | 100 | 49 | 0.880 | 1.040 | 1.180 | 1.040 | 43 | 51 | 58 | 51 |
| 06 | F | 30 | 100 | 56 | 1.000 | 0.740 | 1.230 | 1.200 | 56 | 41 | 69 | 67 |
| 07 | M | 31 | 100 | 39 | 1.100 | 1.030 | 0.940 | 1.050 | 43 | 40 | 37 | 41 |
| 08 | M | 38 | 100 | 45 | 1.620 | 1.150 | 1.190 | 0.930 | 73 | 52 | 53 | 42 |
| 09 | F | 27 | 100 | 39 | 1.460 | 1.180 | 1.330 | 1.220 | 57 | 46 | 52 | 48 |
| 10 | F | 24 | 80 | 40 | 1.660 | 1.200 | 1.340 | 1.380 | 67 | 48 | 54 | 55 |
| 11 | M | 34 | 73 | 47 | 1.410 | 1.010 | 1.240 | 1.020 | 67 | 48 | 58 | 48 |
| 12 | M | 31 | 100 | 59 | 1.340 | 0.810 | 1.400 | 0.910 | 79 | 48 | 83 | 54 |
| 13 | F | 25 | 80 | 50 | 1.020 | 1.020 | 1.300 | 1.170 | 51 | 51 | 65 | 59 |
| 14 | F | 25 | 87 | 57 | 0.930 | 0.900 | 0.860 | 1.020 | 53 | 51 | 49 | 58 |
| 15 | M | 26 | 100 | 56 | 0.820 | 0.770 | 0.980 | 0.810 | 46 | 43 | 55 | 45 |
| 16 | F | 22 | 75 | 56 | 1.300 | 1.280 | 1.420 | 1.330 | 73 | 72 | 80 | 75 |
| 17 | F | 22 | 100 | 44 | 1.120 | 1.160 | 1.250 | 0.840 | 49 | 51 | 55 | 37 |
| 18 | F | 29 | 85 | 32 | 1.260 | 1.480 | 1.420 | 1.400 | 40 | 47 | 45 | 45 |

*Notes:* LQ: Handedness laterality quotient. rMT = resting motor threshold in % MSO (maximum stimulator output). SOM = somatomotor cortex; AUD = auditory cortex; IPL = inferior parietal lobe; DLPFC = dorsolateral prefrontal cortex. SOM Ratio, AUD Ratio, IPL Ratio, and DLPFC Ratio columns provide scaling factors for e-field based dosing with respect to the resting motor threshold (rMT). SOM %, AUD %, IPL %, DLPFC % columns provide the resulting stimulator intensities in % MSO.

**Table S2.** Sample statistics - Group level

| | Age | LQ | rMT | SOM Ratio | AUD Ratio | IPL Ratio | DLPFC Ratio | SOM % | AUD % | IPL % | DLPFC % |
|---|---|---|---|---|---|---|---|---|---|---|---|
| **MEAN** | 30.50 | 90.83 | 46.39 | 1.264 | 1.039 | 1.215 | 1.111 | 57.61 | 47.39 | 56.33 | 51.11 |
| **SD** | 5.88 | 11.01 | 8.15 | 0.268 | 0.255 | 0.285 | 0.237 | 11.58 | 11.00 | 15.66 | 12.06 |
| **MIN** | 22 | 73 | 32 | 0.820 | 0.730 | 0.850 | 0.810 | 40 | 26 | 31 | 33 |
| **MAX** | 40 | 100 | 59 | 1.660 | 1.610 | 2.040 | 1.730 | 79 | 72 | 86 | 75 |

*Notes:* LQ: Handedness laterality quotient. rMT = resting motor threshold in % MSO (maximum stimulator output). SOM = somatomotor cortex; AUD = auditory cortex; IPL = inferior parietal lobe; DLPFC = dorsolateral prefrontal cortex. SOM Ratio, AUD Ratio, IPL Ratio, and DLPFC Ratio columns provide scaling factors for e-field based dosing with respect to the resting motor threshold (rMT). SOM %, AUD %, IPL %, DLPFC % columns provide the resulting stimulator intensities in % MSO.

#### Relationship between cortical stimulation thresholds & modeling fits

In the main text, we argue that exceptionally high cortical stimulation thresholds in some participants (as compared to other participants) may be due to suboptimal modeling, rather than genuinely higher stimulation thresholds in these individuals. This notion is supported by a negative correlation between cortical stimulation thresholds (i.e., the e-field strength in the M1 hotspot at resting MT; V/m) and modeling fits between the e-field magnitude and MEP amplitude at the motor hotspot ($R^2$) (Figure S1). In other words, participants with worse modeling fits show higher cortical stimulation thresholds ($r = -0.56$, $p = 0.019$).

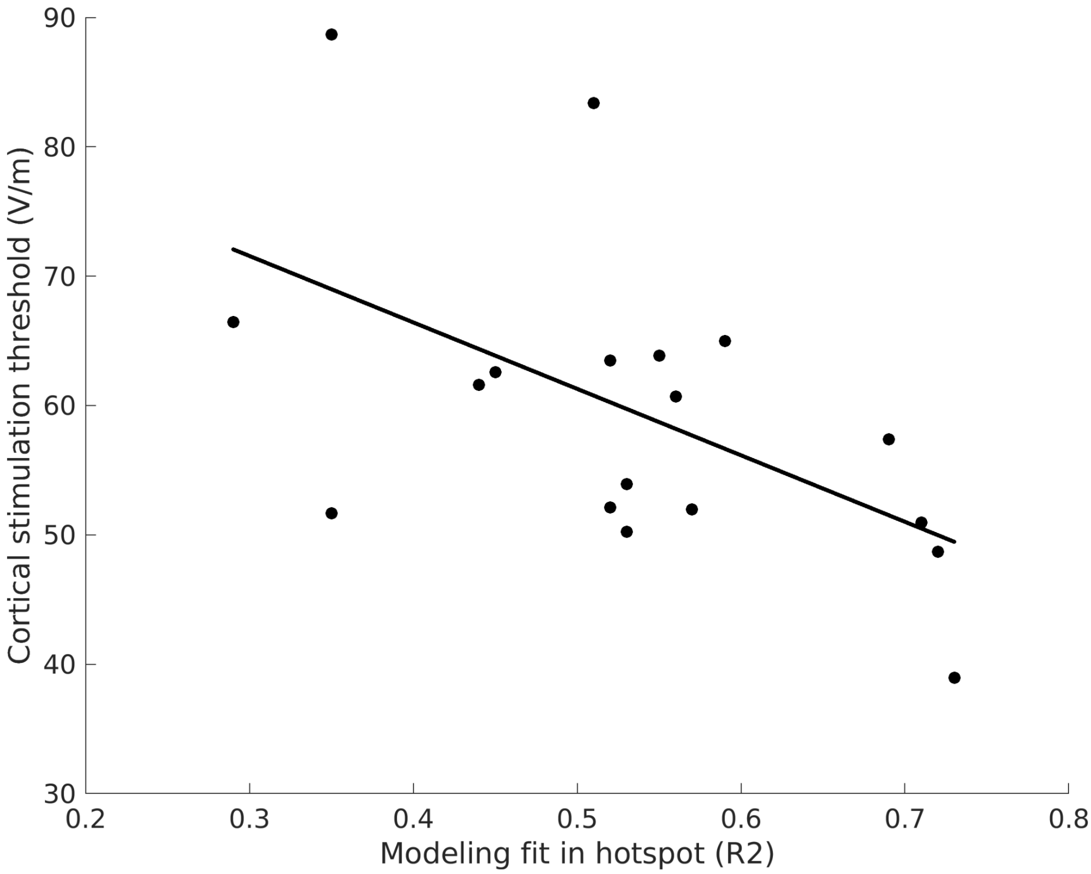


**Figure S1.** Correlation of individual fits between the e-field magnitude and MEP amplitude at the motor hotspot ($R^2$) and cortical stimulation thresholds (V/m).

### References

Numssen, O., Zier, A. L., Thielscher, A., Hartwigsen, G., Knösche, T. R., & Weise, K. (2021). Efficient high-resolution TMS mapping of the human motor cortex by nonlinear regression. *NeuroImage*, *245*, 118654. DOI: 10.1016/j.neuroimage.2021.118654

Puonti, O., Van Leemput, K., Saturnino, G. B., Siebner, H. R., Madsen, K. H., & Thielscher, A. (2020). Accurate and robust whole-head segmentation from magnetic resonance images for individualized head modeling. *NeuroImage*, *219*, 117044. DOI: 10.1016/j.neuroimage.2020.117044

Saturnino, G. B., Puonti, O., Nielsen, J. D., Antonenko, D., Madsen, K. H., & Thielscher, A. (2019). SimNIBS 2.1: A Comprehensive Pipeline for Individualized Electric Field Modelling for Transcranial Brain Stimulation. In: Makarov, S., Horner, M., Noetscher, G. (Eds.), *Brain and Human Body Modeling* (3-25). Springer, Cham. DOI: 10.1007/978-3-030-21293-3_1

Thielscher, A., Antunes, A., & Saturnino, G. B. (2015). Field modeling for transcranial magnetic stimulation: A useful tool to understand the physiological effects of TMS? *Annu. Int. Conf. IEEE Eng. Med. Biol. Soc., 2015,* 222–225. DOI: 10.1109/embc.2015.7318340

Wagner, T. A., Zahn, M., Grodzinsky, A. J., & Pascual-Leone, A. (2004). Three-dimensional head model simulation of transcranial magnetic stimulation. *IEEE Trans. Biomed. Eng.*, *51*(9), 1586–1598. DOI: 10.1109/TBME.2004.827925

Weise, K., Numssen, O., Kalloch, B., Zier, A. L., Thielscher, A., Haueisen, J., Hartwigsen, G., & Knösche, T. R. (2022). Precise motor mapping with transcranial magnetic stimulation. *Nat. protoc.*, *18*(2), 293–318. DOI: 10.1038/s41596-022-00776-6
